# Structure and mechanism of the broad spectrum CRISPR-associated ring nuclease Crn4

**DOI:** 10.1101/2025.07.17.665281

**Authors:** Haotian Chi, Ville Hoikkala, Stephen McMahon, Shirley Graham, Tracey Gloster, Malcolm F White

## Abstract

Type III CRISPR systems detect the presence of RNA from mobile genetic elements (MGE) in prokaryotes, providing antiviral immunity. On activation, the catalytic Cas10 subunit conjugates ATP to form cyclic oligoadenylate (cOA) signalling molecules that activate ancillary effectors, providing an immune response. Cellular ring nucleases degrade cOA to reset the system. Here, we describe the structure and mechanism of a new family of ring nucleases, Crn4, associated with type III-D CRISPR systems. The crystal structure of Crn4 reveals a small homodimeric protein with a fold unrelated to any known ring nuclease or, indeed, any known protein structure. Crn4 degrades a range of cOA species to linear oligoadenylates *in vitro* and ameliorates type III CRISPR immunity *in vivo*. Phage and plasmids also encode Crn4 orthologues that may function as anti-CRISPRs. These observations expand our understanding of ring nucleases and reveal a new protein fold for cyclic nucleotide recognition.

## Introduction

CRISPR-Cas systems provide adaptive immunity against invading mobile genetic elements (MGE) in prokaryotes ^1^. Type III CRISPR-Cas systems comprise a ribonucleoprotein effector complex programmed with CRISPR RNA (crRNA) to detect invading RNA from viruses and other MGE ^2^. Target RNA binding activates the catalytic Cas10 subunit, which in approximately 90% of cases harbours a specialised polymerase (PALM) domain for the synthesis of cyclic oligoadenylate (cOA) molecules ^3–6^. cOA acts as a second messenger, signalling viral infection in the cell by binding and activating a wide range of ancillary defence proteins (reviewed in ^7^). Cleavage of bound target RNA deactivates the Cas10 cyclase activity ^8^ and extant cOA molecules are degraded by enzymes known as ring nucleases ^9^.

The Csm6/Csx1 family of type III CRISPR ancillary effectors use a dimeric CARF domain for cOA binding, resulting in allosteric activation of their HEPN-family ribonuclease domains ^5,6^. Some of these e\ectors also degrade cA4 or cA6 in their CARF domains, providing a mechanism for auto-deactivation of the CRISPR immune response ^10–14^. The effectors Cami1 and CalpL, which detect cA4 via CARF and SAVED domains, respectively, also possess intrinsic ring nuclease activity ^15,16, 17^.

The first extrinsic ring nuclease to be described, Crn1, was purified from *Saccharolobus solfataricus*, revealing a small protein with a dimeric CARF domain for cOA recognition. Crn1 slowly degrades cyclic tetra-adenylate (cA4) to linear di-adenylate products ^18^. A second family of extrinsic ring nucleases, Crn2, use a dimeric domain unrelated to CARF proteins to bind and degrade cA4. Crn2 is found both in CRISPR defence operons ^19–21^ and in viral genomes, where it functions as an anti-CRISPR (Acr) by rapidly degrading cA4 to neutralise cellular defences ^21^. A third ring nuclease family, Crn3, uses a fold related to CARF domains to sandwich cA4 in protein tetramers, degrading the signalling molecule in a metal-dependent reaction to linear di-adenylates ^22,23^. Finally, a group of related proteins (Csx15, Csx16 and Csx20) were predicted to act as ring nucleases ^24^. Recently, we demonstrated that Csx16 and Csx20 are cA4-specific ring nucleases, and showed that the majority of cA4- and cA6-signalling type III CRISPR systems include a means to degrade their activator ^25^. This has important implications for cellular outcomes on viral infection, as it provides a means to avoid programmed cell death or growth arrest ^25,26^.

We previously reported a systematic bioinformatic analysis of type III CRISPR systems which resulted in the identification of a new effector protein, Csm6-2, which is activated by cyclic hexa-adenylate (cA6) and cleaves RNA non-specifically ^3^. Csm6-2 is typically found associated with a CorA-family trans-membrane effector, and in some instances the two proteins are fused. The CRISPR-associated CorA effector from *Bacteroides fragilis* was previously shown to be activated by SAM-AMP – a signalling molecule derived from conjugation of ATP and S-adenosyl methionine ^27^, but systems encoding both Csm6-2 and CorA may signal via cA6 ^3^.

Here, we show that these CRISPR loci include a small open reading frame encoding a hypothetical protein, hereafter named Crn4 (CRISPR ring nuclease 4), which degrades cA3, cA4 and cA6 to a range of linear products. The crystal structure of Crn4 reveals a novel, dimeric fold with a conserved histidine residue that is essential for catalysis. Crn4 can partially relieve CRISPR-mediated immunity in a plasmid challenge assay, consistent with a function as a cellular ring nuclease, and is encoded in some phage and plasmid genomes where it likely functions as an Acr.

## Results

### Identification of the CRISPR associated protein Crn4

Csm6-2, a fused Csm6 orthologue that is activated by cA6 and cleaves RNA, was recently identified from a bioinformatic analysis of type III CRISPR loci ^3^. A systematic scan of the regions neighbouring Csm6-2 revealed a small open reading frame (ORF) encoding a hypothetical protein of unknown function in 12 CRISPR type III-D loci. In 5 of the 12 instances, a CorA membrane effector was also present (Figure 1A). In two cases, the ORF was fused with Csm6-2, and in one instance it was fused with the effector TIR-SAVED (Figure 1B) ^3,28^. A recent bioinformatic study identified this ORF, named Unk01, as a potential ring nuclease ^24^.

**Figure 1.**
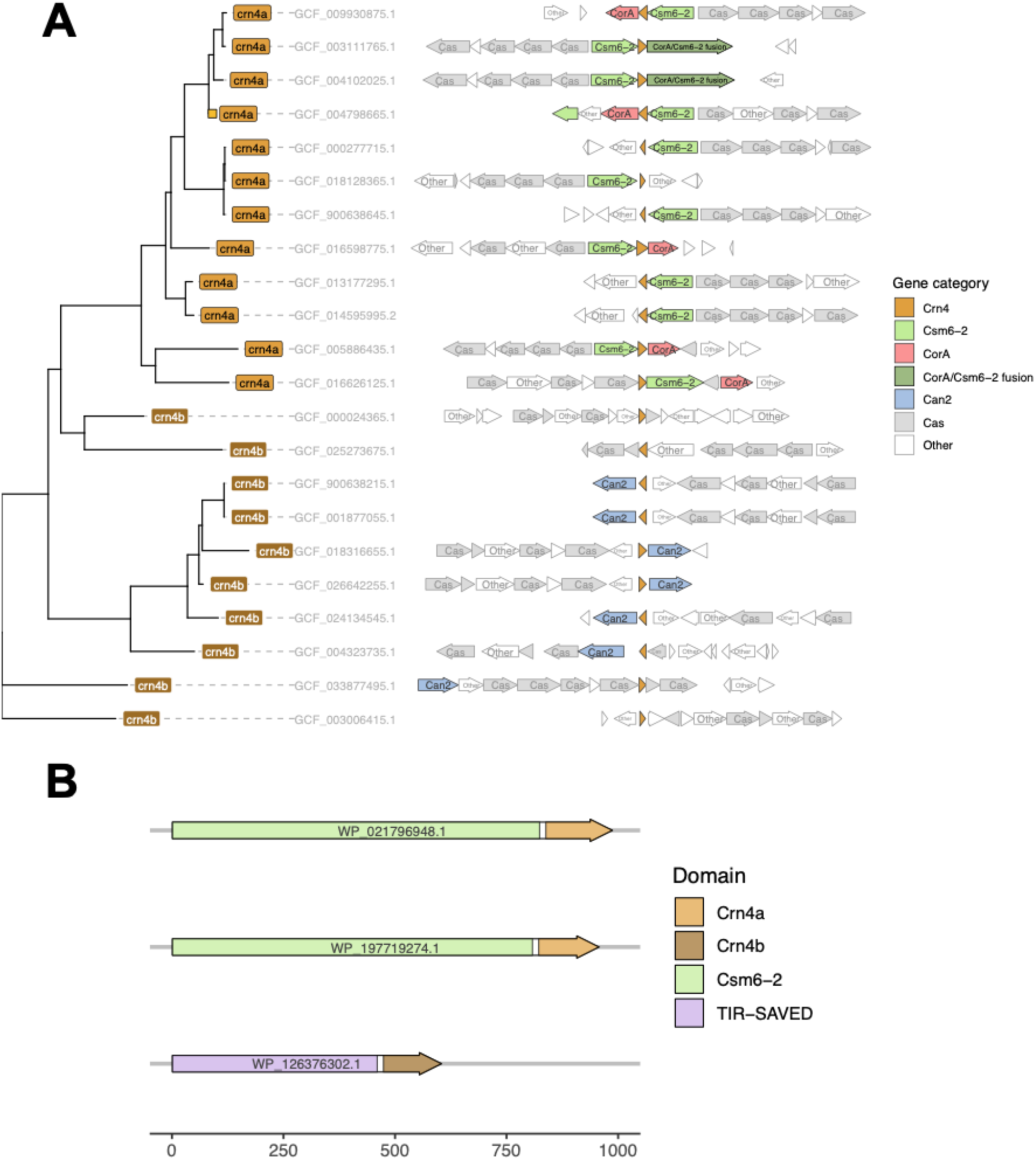
Bioinformatic Analysis of Crn4. **A.** A phylogenetic tree of Crn4 homologues found in type III CRISPR-Cas loci of complete prokaryotic genomes in NCBI. The subfamily of each Crn4, a or b, is labelled on the end nodes followed by NCBI genomic accession numbers. The closest homologue (94% similarity) of the Crn4a tested experimentally in this study is marked by a small square. The genomic neighbourhoods of each Crn4, defined by CRISPR-Cas locus boundaries, is shown on the right, with key e\ectors highlighted as shown in the legend. **B.** Three examples of Crn4 fused to the C-terminus of type III CRISPR-Cas e\ector proteins Csm6-2 and TIR-SAVED. Protein accession numbers are shown on the annotations.

Using hidden Markov model (HMM) profiles based on these 12 homologues, we identified 10 additional homologues in our dataset, which were associated with the cA4-activated e\ector Can2, and two homologues with no recognisable effector in the CRISPR locus. Phylogenetic analysis divided the 22 homologues into two distinct groups: group “a” associated with Csm6- 2/CorA, and group “b” associated with Can2 ^29,30^ or lacking a recognizable effector (Figure 1A). The small size of the predicted proteins, combined with their consistent association with cA4 and cA6 activated e\ectors (either through adjacency or fusion) and prevalence on MGEs supported the prediction that Unk01 was a ring nuclease. Following established nomenclature, we provisionally designated this family of proteins CRISPR ring nuclease 4 (Crn4). The family was sub-divided into Crn4a and Crn4b, distinguishable as Crn4b members are smaller, lacking a ∼10 amino acid region present in Crn4a (Supplementary Figure 1).

Crn4b is also observed in the genomes of phages and plasmids. Of the 28,137 phage genomes in the Millard database ^31^, 11 contained a Crn4 homologue, while this was true for 10 out of 59,895 plasmid sequences in the database PLSDB ^32^. Some of the phage and plasmid orthologues have an 60-80 amino acid N-terminal extension of unknown function (Supplementary Figure 2). All members of the family have conserved residues including T12, H15 and R112 (*A. procaprae* Crn4a numbering), which are implicated in cOA binding and catalysis (further details below).

### Crn4a is a ring nuclease with broad specificity

We chose candidate Crn4a protein WP_136192672, found in the *Actinomyces procaprae* type III CRISPR locus adjacent to the previously characterised Csm6-2 protein ^3^, for further study. To explore its function, we constructed a synthetic gene encoding the protein for expression in *E. coli* and purified it by immobilised metal affinity and size exclusion chromatography (Supplementary Figure 3). Crn4a was tested for ring nuclease activity by incubating the purified protein with synthetic cOA species under multiple turnover (substrate excess) conditions in EDTA, followed by HPLC analysis (Figure 2A). Crn4a cleaved all three cOA species, cA6, cA4 and cA3, generating linear products. cA6 was degraded with a rate constant of 0.024 ± 0.007 min^-1^ (Figure 2B; Supplementary Figure 4), cA3 slightly faster (0.083 ± 0.02 min^-1^) and cA4 was degraded much more rapidly (0.98 ± 0.1 min^-1^). cA4 and cA6 were cleaved on both sides of the ring, yielding linear 2’,3’-cyclic phosphate cA2>P and cA3>P products, respectively (Figure 2A; Supplementary Figure 4). This suggests concerted or sequential cleavage of cOA on opposite sides of the molecules, reminiscent of the viral ring nuclease AcrIII-1 ^21^. cA3, which unlike cA4 and cA6 lacks an axis of two-fold symmetry, was degraded to A2>P and A1>P, consistent with asymmetric cleavage (Figure 2A; Supplementary Figure 4). The ability to cleave multiple cOA species was not observed for Crn1-3 ^18,21,22^. Thus, in contrast to all other CRISPR associated ring nucleases tested to date, Crn4a has broad specificity for cOA species and is the first confirmed CRISPR-associated cA3 ring nuclease.

**Figure 2.**
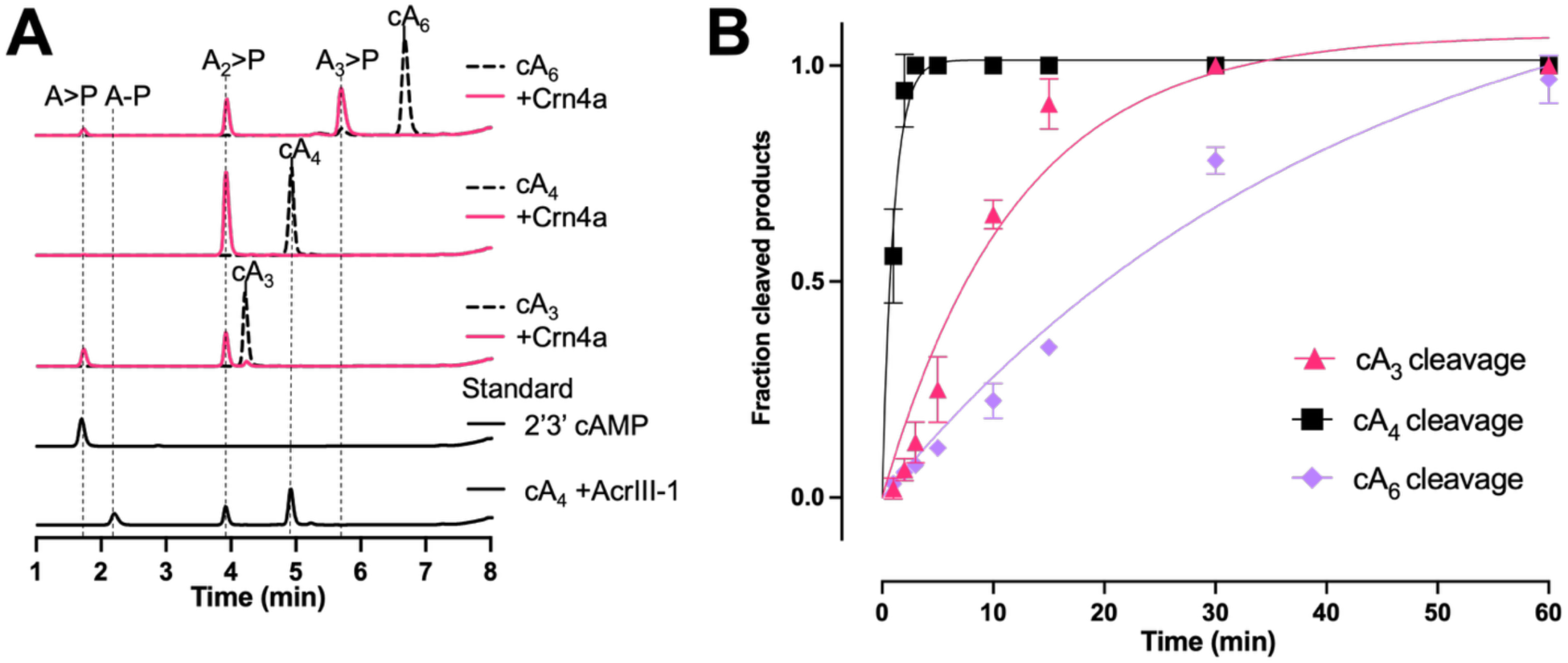
Cleavage of cOA species by Crn4a. **A.** HPLC analysis of cleavage products of cA3, cA4 and cA6 incubated with *A. procaprae* Crn4a. 2’,3’-cAMP and a control reaction cleaving cA4 with AcrIII-1 to generate A2>P are shown as standards. **B.** Kinetic analysis of ring nuclease activity of Crn4a against cA3, cA4 and cA6. Following HPLC, substrate and product peaks were quantified and data plotted as fraction cleaved against time. Data points represent the means of triplicate experiments and standard deviation is shown.

We also cloned and expressed the gene encoding a representative of the Crn4b family (MBK8772583 from *Hyphomicrobiales bacterium*) (Supplementary Figures 1, 2). Crn4b degraded cA4 and cA3 but was inactive against cA6 (Figure 3A). This appears consistent with the association of Crn4b with the Can2 effector, which is activated by cA4 ^29,30,33^. Although Crn4b produces the same final cA4 cleavage products as Crn4a, intermediate linear products (A3>P and A4>P) were observed at early reaction time points with cA3 and cA4, respectively (Figure 3A; Supplementary Figure 5), consistent with a sequential cleavage mechanism.

**Figure 3.**
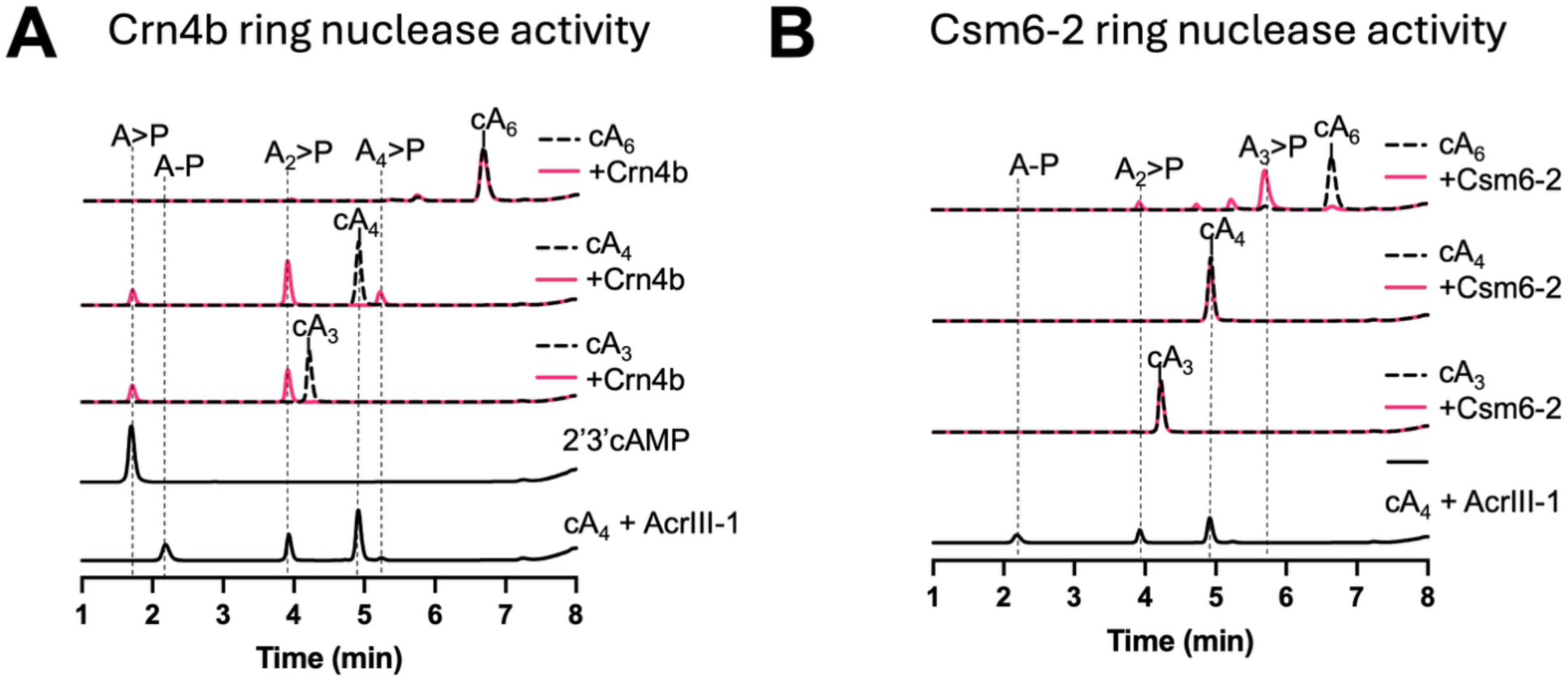
Ring nuclease activity of Crn4b and Csm6-2. **A.** HPLC analysis of cleavage products of cA3, cA4 and cA6 incubated with Crn4b. Controls are as for figure 2A. **B.** *A. procaprae* Csm6-2 possesses ring nuclease activity against cA6, but not cA3 or cA4.

The close association of Crn4a with the Csm6-2 effector prompted us to test the ring nuclease activity of Csm6-2, as all Csm6 proteins studied to date possess intrinsic cA6 ring nuclease activities in the CARF domain of the protein ^10,11,13^. Assays carried out in EDTA, to inhibit the metal ion dependent RNase activity of the HEPN domain, revealed intrinsic cA6 ring nuclease activity for Csm6-2 (Figure 3B). The rate of cleavage of cA6, 0.026 ± 0.01 min^-1^ (Supplementary Figure 6), was equivalent to that observed for Crn4a. Neither cA3 nor cA4 were degraded by Csm6-2, however.

### Structure and mechanism of Crn4

To characterise the structure of Crn4, we crystallised the wild-type Crn4b protein in apo-form, and the Crn4a H15A variant in the presence of cA6 and collected X-ray diffraction data to 1.09 and 1.44 Å resolution, respectively. The structures were solved using molecular replacement with the AlphaFold predicted structure as the model. The structures reveal a new protein fold, with no hits identified by the DALI server ^34^ with Z-scores higher than 2.6. The structure comprises four (Crn4b) or five (Crn4a) pairs of beta strands, one parallel and the others anti- parallel, and one short alpha helix. Both proteins are dimeric, and strikingly, an extended β- hairpin from one monomer wraps around the neighbouring subunit which is largely facilitated by hydrophobic interactions between the two molecules (Figure 4A). The structures of apo Crn4b and Crn4a bound to cA6 superimpose with an RSMD of 2.7 Å over 96 Cα residues (Figure 4B). Whilst they show a similar arrangement for the globular part of the protein, the extended β-hairpin bends at an “elbow” in Crn4a, compared to Crn4b, thus adopting a different conformation with the β-hairpin approximately perpendicular to each other. In addition, the loop of the hairpin is around 10 residues longer in Crn4a. Conserved residues His-15 and Arg-112 (Crn4a numbering) are situated on loops that sit above the central cavity in suitable positions to participate in cOA binding and/or cleavage (Figure 4A). The Crn4a dimer binds its cA6 ligand in a symmetrical arrangement, with two adenine bases oriented upwards, two downwards and two roughly axial to the plane of the cA6 ring (Figure 4C,D). This differs from the conformation of cA6 bound to the ribonuclease *Streptococcus thermophilus* Csm6, where four of the six adenine bases were in the plane of the ring ^35^. In Crn4a, cA6 hydrogen bonds with the main chain atoms of Arg92, Glu16, Pro77, Ala70, and side chain hydroxyl of Ser91, and a phosphate moiety forms an electrostatic interaction with Arg112. In addition, there is pi-pi stacking between the adenine of cA6 and guanidium group of the Arg92. Arg92, Glu16, Ser91 and Arg112 are all structurally conserved in Crn4b, as are the main chain atoms of Pro77 and Ala70 but with alternative side chains. Both Arg92 and Arg112 are present in two conformations in Crn4a, and whilst conserved are found in a different orientation in Crn4b, indicating these residues have a fair amount of flexibility and likely to stabilise upon binding of cA6. The orientation of Arg112, and its interaction with a phosphate group in cA6 points to this being an important residue in phosphodiester bond cleavage. In addition, alignment of the Crn4a H15A mutant with Crn4b suggests His15 is in a key position to participate in catalysis, with Thr13 positioned just behind it (Figure 4E).

**Figure 4.**
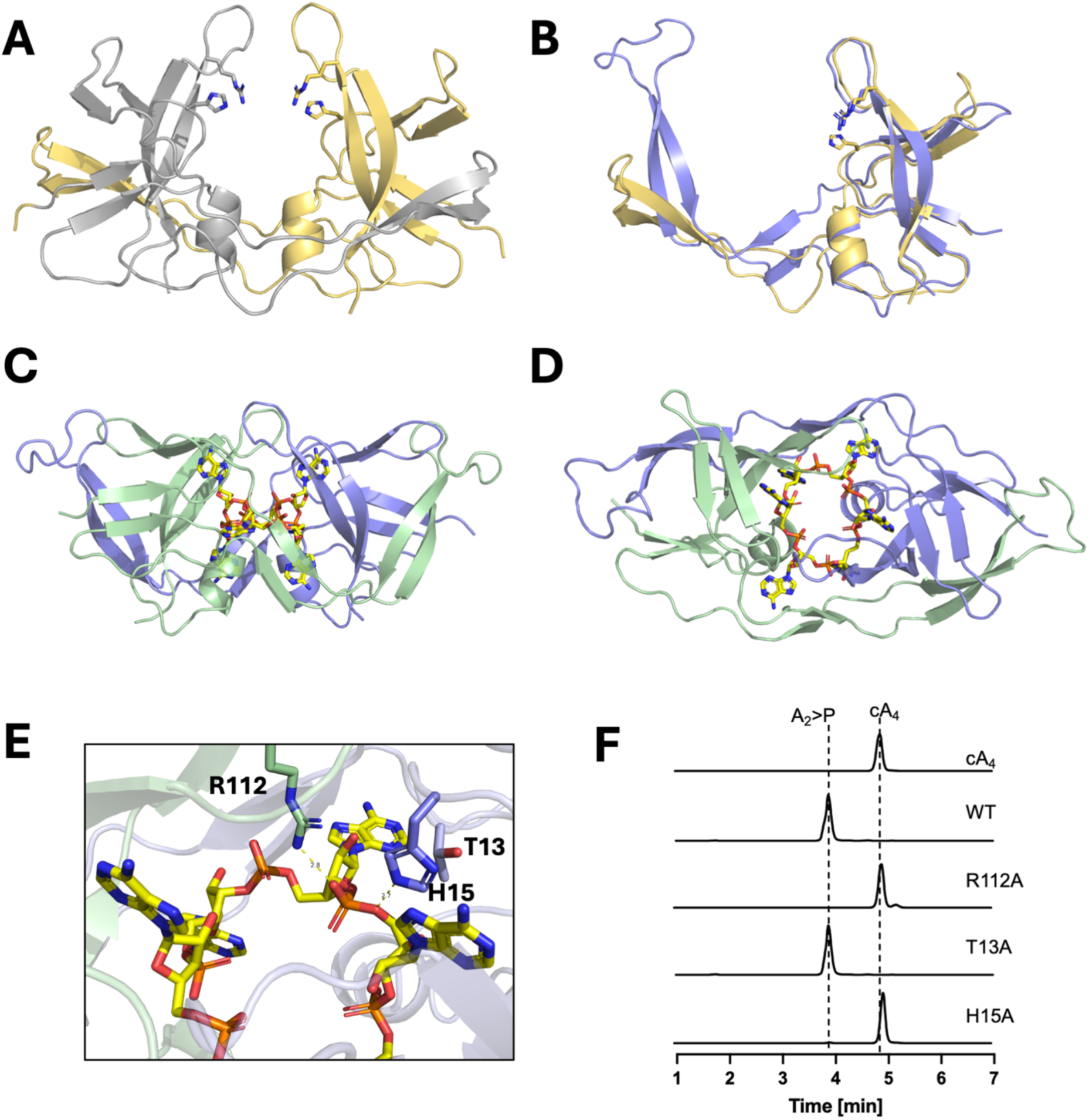
Structure and mechanism of Crn4. **A.** Dimeric structure of Crn4b, with subunits coloured grey and gold. Each subunit has an extended β-hairpin arm around the other. The conserved residues His17 and Arg103 are positioned on loops above a central cavity. **B.** Superimposition of a monomer of Crn4b (gold) and Crn4a (blue). Crn4a has a more extensive and bent β-hairpin arm compared to Crn4b. The conserved residues His17 and Arg103 (Crnb4b) and Ala15 (mutated from histidine) and Arg112 (Crn4a) are shown. **C.** Dimeric structure of Crn4a, with subunits coloured blue and green. cA6 bound to Crn4a is shown in stick representation, with carbons coloured yellow. **D.** Top-down view of the Crn4a dimer, highlighting how the extended β-hairpin wraps around the neighbouring subunit. **E.** Active site of Crn4a, with subunits coloured green and blue. Conserved residues His15 and Arg112 are suitably positioned to participate in cA6 binding and turnover, with Thr13 sitting behind His15. His15 is modelled into the H15A Crn4a structure, guided by the position and orientation of the equivalent residue observed in the Crn4b structure **F.** Mutagenesis of His15 and Arg112 abolished ring nuclease activity, while the T13A variant was still active.

We constructed and purified variants T13A, H15A and R112A of Crn4a to test their role in catalysis. Mutation of His15 or Arg112 to alanine abolished cleavage of the preferred substrate, cA4, after incubation at 37 °C for 60 min, while the T13A variant had wild-type levels of activity (Figure 4F). Qualitatively similar data were obtained for cA3 and cA6 cleavage (Supplementary Figure 7). These data are consistent with a role for His15 as a general acid, protonating the oxyanion leaving group during phosphodiester bond cleavage, reminiscent of His47 in AcrIII-1 ^21^. Arg112 from the neighbouring subunit likely provides a key electrostatic interaction partner for the bound cOA species during binding and catalysis.

### Crn4 is functional in vivo

To explore a functional role of Crn4 *in vivo*, we reconstituted *A. procaprae* Crn4a into a well-developed recombinant type IIIA CRISPR system from *Mycobacterium tuberculosis* (MtbCsm), which produces a range of cOA species for effective activation of cOA-dependent effectors to provide plasmid immunity ^36^. *E. coli* C43 cells expressing MtbCsm were transformed with a pRATDuet plasmid expressing an effector protein alone or along with Crn4a and carrying a tetracycline resistance gene (*tetR*). Fewer transformants were expected when the effector was activated by cOA generated by the MtbCsm system, which was programmed with a CRISPR RNA (crRNA) targeting a portion of *tetR* gene. As a control, we also tested MtbCsm programmed with a crRNA targeting the pUC plasmid, which is not activated to generate cOA in this experimental setup. Effectors tested in this programmed MtbCsm system were cA4-activated Csx1 from *Thioalkalivibrio sulfidiphilus* (TsuCsx1) ^27,37^ and cA6- activated e\ectors Csm6 from *M. tuberculosis* (MtbCsm6) ^36^ and *A. procaprae* Csm6-2 ^3^.

In the absence of Crn4a, all three effectors were activated on transformation of the pRATDuet plasmid, providing effective resistance to plasmid transformation, reflected in reduced colony forming units (cfu) (Figure 5; Supplementary figure 8). The presence of Crn4a effectively alleviated plasmid immunity conferred by cOA-activated TsuCsx1 and MtbCsm6 (Figure 5), suggesting that Crn4a reduces the concentration of both cA4 and cA6 when expressed in cells. In contrast, no significant difference in colony counts was observed for the Csm6-2 effector in the presence or absence of Crn4a. This may be because Csm6-2 is itself a very active ring nuclease; alternatively this could reflect the relative a\inities of cA6 binding by Crn4a and Csm6-2.

**Figure 5.**
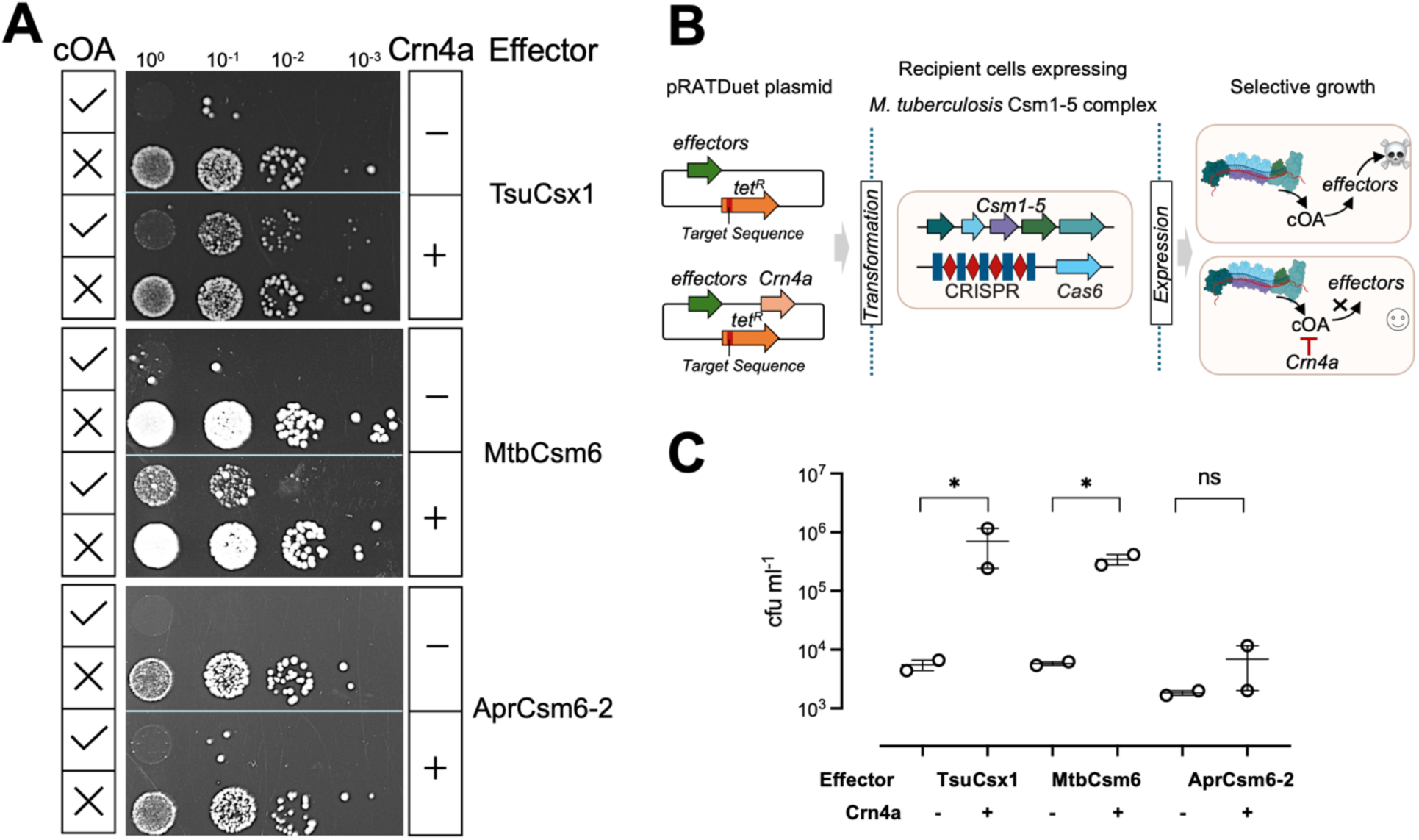
Crn4a neutralises CRISPR immunity in a plasmid challenge assay. **A.** Plasmid challenge assay. The expression of Crn4a increased cfu in cells with cA4-activated TsuCsx1 and cA6-activated MtbCsm6, consistent with degradation of those cOA species. For the Csm6-2 effector however, no increase in cfu was observed when Crn4a was co-expressed. cOA was synthesised in the MtbCsm system programmed with crRNA targeting the *tetR* gene, while no cOA production in the MtbCsm non-target (pUC) system was set as a negative control (represented by ticks and crosses, respectively). Representative plates of two biological replicates with two technical replicates each are shown. **B.** Schematic of the experimental design. **C.** Colony counts for transformants in the presence or absence of Crn4a in the context of activated MtbCsm target system. Data are presented as mean ± s.d. Statistical analysis was performed using ratio paired t test (one-tailed *P* values listed from left to right on the graph: \**P*= 0.0399, \**P*= 0.0217 and ^ns^*P*= 0.25).

## Discussion

Ring nucleases, first described in 2018 ^18^, are now understood to be an intrinsic aspect of type III CRISPR defence, where it seems that an ability to degrade the cOA second messengers generated by these systems is generally beneficial to the host ^25^. Of the six types of specialised or stand-alone ring nucleases characterised to date, all are based on Rossmannoid domain architecture (eg. CARF and SAVED domains) ^25^. Furthermore, all of them are specific for degradation of cA4, the most common cOA signalling species ^3^.

Meanwhile, the self-limiting effectors with intrinsic ring nuclease activity also use CARF or SAVED domains to degrade either cA6 or cA4 ^10–17^, although the CalpL protein also has some activity against cA3 *in vitro* ^16^ . Here, we have described a new family of ring nucleases, Crn4, with several notable characteristics. Firstly, Crn4 displays a novel (non-Rossmann) protein fold for cOA binding. Secondly, Crn4 has ring nuclease activity against all the cOA species known to signal in type III CRISPR defence.

The structure of Crn4 appears to be completely unique when compared to others in the Protein Databank, thus expanding our understanding of the protein folds that can bind cyclic nucleotides. The chemistry of cOA cleavage, on the other hand, is highly reminiscent of previously characterised ring nucleases, such as AcrIII-1, with both enzymes positioning an essential histidine as a likely general acid to facilitate catalysis ^21^. Cleavage of these nucleotide species is relatively straightforward once the cOA molecule is bound in the correct orientation, allowing the 2’-hydroxyl of the ribose sugar to attack the phosphodiester bond, so there seems to be little obstacle to ring nuclease activity evolving in protein domains that bind cOA species.

The detection of Crn4b homologues in phage and plasmid genomes implies a role for these enzymes as anti-CRISPR proteins, analogous to the cA4-specific AcrIII-1 ^21^. The activity of Crn4b against both cA3 and cA4 suggests that both signalling molecules could be intercepted by these Acrs, although that remains to be confirmed. Viral nucleases targeting cA3 could potentially function as anti-defence enzymes against both type III CRISPR and CBASS (cyclic nucleotide based anti-phage signalling systems), each of which utilise this signalling molecule ^38–41^. To date, only the broad specificity anti-defence phosphodiesterase Acb1 has been shown capable of degrading cA3 ^42^.

We first identified Crn4 as a focus for investigation based on its close association with the cA6-activated CRISPR effector Csm6-2 ^3^. As we have shown, Csm6-2 has intrinsic ring nuclease activity in its CARF domains and reaction rates *in vitro* are comparable to those for Crn4a, raising the question of why the latter protein is required? In fact, effectors with intrinsic ring nuclease activity are commonly found in type III CRISPR loci alongside dedicated ring nucleases ^25^, suggesting there is an advantage to the host organism to have such an arrangement. One possibility is that the relatively high levels of cOA species generated by an activated type III CRISPR system ^8^ necessitate a mechanism to remove these molecules when their function is no longer needed – one that does not rely on the activation, however transient, of a toxic effector protein.

In summary, these findings revealed a new class of cyclic nucleotide binding proteins, highlighting the wide distribution and diversity of ring nucleases in type III CRISPR defence.

## Methods

### Cloning

The g-blocks of the *crn4a* and *crn4b* genes were codon optimised for expression in *E. coli* and purchased from IDT (Integrated DNA Technologies). Synthetic genes were cloned into the vector pEhisV5TEV ^8^ between the *Nco*I and *BamH*I sites. The *E. coli* DH5a strain was used for cloning, and sequence integrity was confirmed by sequencing (Eurofins Genomics). Site-directed mutagenesis was performed using primers with the desired mutations. All synthetic genes and primers used in this study are listed in Supplementary Table 1. For the plasmid challenge assay, the g-block of the *crn4a* gene was cloned into the MCS-1 (Multiple Cloning Site-1) of pRATDuet ^36^ between the *Nde*I and *Xho*I sites, under the control of a T7 promoter. For the expression of two proteins, the synthetic gene encoding TsuCsx1 ^36^, MtbCsm6 ^36^ or Csm6-2C ^3^ was cloned into the MCS-2 (Multiple Cloning Site-2) of pRATDuet-Crn4 plasmid between the *Nco*I and *Hind*III sites, under the control of pBAD promoter.

### Protein expression and purification

*E. coli* C43 (DE3) cells were transformed with plasmids for protein expression. 1 L of Luria-Broth (LB) containing cells transformed with the plasmid of interest was grown at 37 °C to an OD600 of 0.6-0.8. Crn4a or Crn4b expression was induced with 0.4 mM isopropyl b-D-1- thiogalactopyranoside (IPTG) for 4 hours at 25 °C and 37 °C, respectively. Cell pellets were harvested for purification by centrifugation at 4,000 rpm (Beckman Coulter Avanti JXN-26; JLA8.1 rotor) at 4 °C for 15 min.

The purification procedure was performed using immobilised metal affinity chromatography (IMAC) and size exclusion chromatography, as described previously ^8^. The identity and purity of proteins were verified using SDS-PAGE and the pure protein was aliquoted and stored at -70 °C.

### Ring nuclease activity

To examine the ring nuclease activity, enzymes were incubated with 200-fold molar excess of cOA species in 20 mM Tris-HCl, pH 7.5, 250 mM NaCl and 1 mM EDTA at 37 °C for the time indicated. For calculation of rate constants of wild type Crn4a/b and Csm6-2, 500 nM enzyme was incubated with 100 mM cA3, cA4 or cA6, at 37 °C for the time points 1, 2, 3, 5, 10, 15, 30 and 60 min. Reaction samples were quenched by mixing with two equivalent volumes of cold methanol and vortexed for 30 s, before centrifugation at 4 °C for 15 min. The supernatant was transferred to a new tube and vacuum dried, before resuspension in H2O for HPLC or LC-MS analysis. Substrate and cleaved products were quantified from triplicate measurements and data were fitted using a one phase exponential model (Y=Ymax*(1-e^-k*X^)) using Prism (Version 10.2.2 (341)).

### HPLC and LC-MS analysis

HPLC analysis was conducted on a UltiMate3000 UHPLC system (Thermo Fisher Scientific) equipped with an Accucore™ C18 column (Thermo Fisher Scientific 2.1 x 100 mm, particle size 2.6 µM) for the time-course samples and a C18 column (Kinetex EVO 2.1 x 50 mm, particle size 2.6 µM) for other analyses. The absorbance was monitored at 260 nm and the column temperature was set at 40 °C. Solvent A was 20 mM ammonium acetate, pH 8.5 and B was methanol. The flow rate was set at 0.4 ml/min for the Accucore™ C18 column and 0.3 ml/min for the EVO C18 column. Gradient elution followed the same procedure: 0-0.5 min, 1% B; 0.5- 6 min, 1-15% B; 6-7 min, 100% B. LC-MS analysis on a Eksigent 400 LC coupled to Sciex 6600 QTof mass spectrometer system has been described previously ^27^.

### Protein crystallisation

Crn4a H15A, at 11 mg mL^−1^, was incubated at room temperature for 30 min with 1.2 molar excess of cA6. Crn4b was used at a concentration of 15 mg mL^-1^. Immediately prior to crystallisation both proteins were centrifuged at 14000*g*. Sitting drop vapour diffusion experiments were set up at the nanoliter scale using commercially available crystallisation screens and incubated at 293 K. Optimised Crn4a H15A with cA6 crystals were grown from 23.8% PEG 3350, 0.2M magnesium chloride, and 0.1M Bis-Tris pH 5.5 and apo Crn4b crystals were grown from 20% PEG 6000, 10% ethylene glycol, and 0.1 M calcium chloride. Crystals were cryoprotected with 25% glycerol prior to harvesting and cryo-cooling in liquid nitrogen.

### X-ray data collection, structure solution, and refinement

X-ray data were collected at a wavelength of 0.9537 Å, 100 K, on beamline I04 at the Diamond Light Source, to 1.44 Å (Crn4a H15A) and 1.09 Å (Crn4b) resolution. Data were automatically processed using Xia2 ^43^. The structure was solved by phasing the data using PhaserMR ^44^ in the CCP4 suite ^45^, using a model generated by AlphaFold 3 ^46^, with initial B-factors modelled in Phenix ^47^. Model refinement was achieved by iterative cycles of REFMAC5 ^48^ with manual model manipulation in COOT ^49^. For the Crn4a H15A with cA6 data, electron density for cA6 was clearly visible in the maximum likelihood/σA weighted *F*obs-*F*calc electron density map at 3σ. The coordinates for cA6 were generated in ChemDraw (Perkin Elmer) and the library was generated using Acedrg ^50^, before fitting the molecule in COOT. The quality of each structure was monitored throughout using Molprobity ^51^. Data and refinement statistics are shown in Supplementary Table 2. The coordinates and data have been validated and deposited in the Protein Data Bank with deposition codes 9QS9 for Crn4a H15A, and 9R7B for Crn4b.

### Plasmid challenge assay

Plasmids from the programmed type III MtbCsm system have been described previously, including pCsm1-5 (containing Csm interference genes *cas10*, *csm3*, *csm4* and *csm5* from *M. tuberculosis* and *csm2* from *M. canettii*) ^36^, pCRISPR-TetR and pCRISPR-pUC (containing *M. tuberculosis cas6* and a CRISPR array targeting a tetracycline-resistance gene or a pUC multiple cloning site, respectively) ^36^ and pRATDuet-effectors (carrying tetracycline-resistance gene and *M. tuberculosis* (Mtb) *csm6 or Thioalkalivibrio sulfidiphilus* (Tsu) *csx1*) ^52^.

*E. coli* C43 (DE3) cells carrying plasmids pCsm1-5 and pCRISPR-TetR or pCRISPR-pUC were transformed with 100 ng of pRATDuet derived plasmids containing di\erent e\ectors with or without a Crn4 ring nuclease. After recovering at 37 °C for 2 h, a tenfold serial dilution of cells was applied onto LB agar supplemented with 100 µg/ml ampicillin and 50 µg/ml spectinomycin for determination of cell density of recipients, onto LB agar containing additional 12.5 g/ml tetracycline for determination of transformation efficiency and LB agar with additional 0.2 % (w/v) D-lactose and 0.2 % (w/v) L-arabinose to determine the plasmid immunity mediated by cOA-dependent effectors. Plates were incubated at 37°C overnight, before taking images of plates. Technical duplicates of two biological replicates were performed. Colonies were counted to calculate colony forming units (cfu/ml) and analysed using an unpaired t test to compare the difference between systems in the presence or absence of Crn4 using Prism (Version 10.2.2).

### Bioinformatic analyses

To search for Crn4 homologues in prokaryotes, we downloaded the 38,742 complete prokaryotic genomes from NCBI on March 25^th^ 2024. For phage genomes we used the Millard database of 28,114 genomes (May 2024 release) ^31^ and for plasmids the PLSDB database containing 59,895 sequences ^32^. Based on our initial identification of two main clades for Crn4 homologues, separate Hidden Markov Model (HMM) databases were created for Crn4a and Crn4b. These profiles were integrated into a previously published Snakemake pipeline developed for finding type III CRISPR-Cas loci, effectors and ring nucleases in prokaryotic and in phage genomes ^3^. In short, the pipeline annotates type III CRISPR-Cas loci and searches them for ring nucleases based on HMM profiles and protein length limitations. A separate pipeline was created for examining plasmid sequences.

To create the phylogenetic tree (Figure 1), prokaryotic Crn4a and Crn4b amino acid sequences were concatenated into one file, and aligned using Muscle ^53^. A phylogenetic tree based on the alignment was created with FastTree ^54^ using -wag and -gamma arguments. Two Crn4 homologues (genomes GCF_002285795.1 and GCF_002863905.1) were removed from the dataset due to missing cyclase domains in the accompanying Cas10. The tree was plotted and annotated in RStudio v. XX with ggtree ^55^. The accompanying genomic neighbourhood visualisation was created by merging information from several annotation files output by the pipeline (CCTyper, cATyper, RN_typer) into the original NCBI general features files (.g\). The enhanced annotations were visualised using ggtree’s geom_facet wrapper for gggenes and centered around Crn4. Custom rules were used to bin the genes into the categories shown in Figure 1. Crn4 fusions with known effector proteins were discovered by modifying the snakemake pipeline to search for Crn4 without protein length limitations.

### Data availability

The protein structure coordinates and data have been deposited in the Protein Data Bank with deposition codes 9QS9 for Crn4a and 9R7B for Crn4b.

## Acknowledgements

Thanks to Dr Sabine Grüschow for helpful discussion. This work was supported by a European Research Council Advanced Grant (Grant REF 101018608 to MFW).

## Author contributions

HC: data acquisition and interpretation. VH: bioinformatic data acquisition and analysis. SM: crystallographic data acquisition and interpretation. SG: data acquisition. TM & MFW: funding acquisition, conception, data interpretation. All authors contributed to the drafting and revision of the manuscript.

---

**Supplementary Figure 1.**
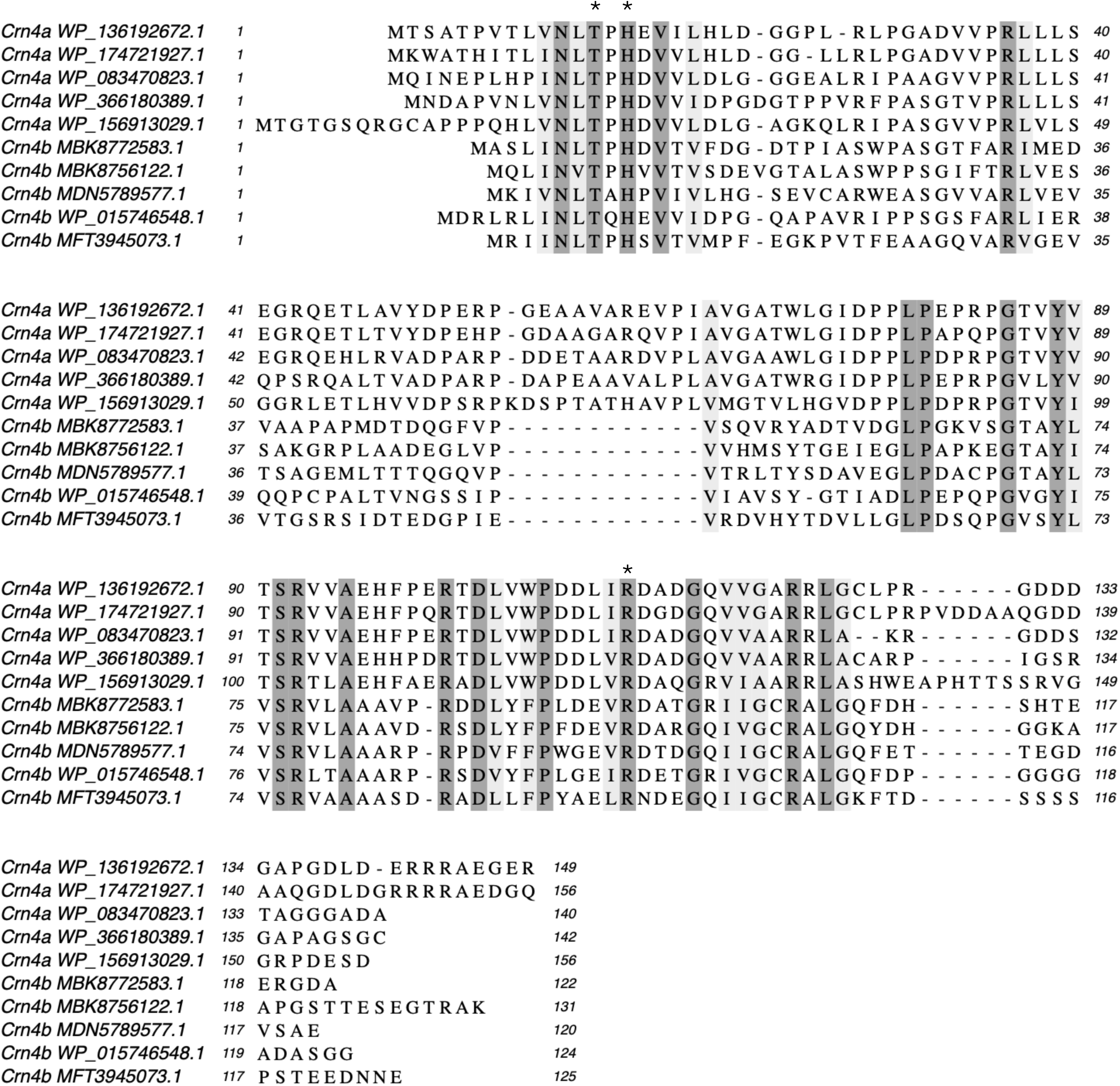
Multiple sequence alignment of a selection of Crn4a and Crn4b proteins. *A. procaprae* Crn4a (WP_136192672.1) and Crn4b (MBK8772583.1) are studied biochemically in this manuscript. Conserved residues are shaded. T12, H15 and R112 (AprCrn4a numbering) are highlighted with asterisks.

**Supplementary figure 2.**
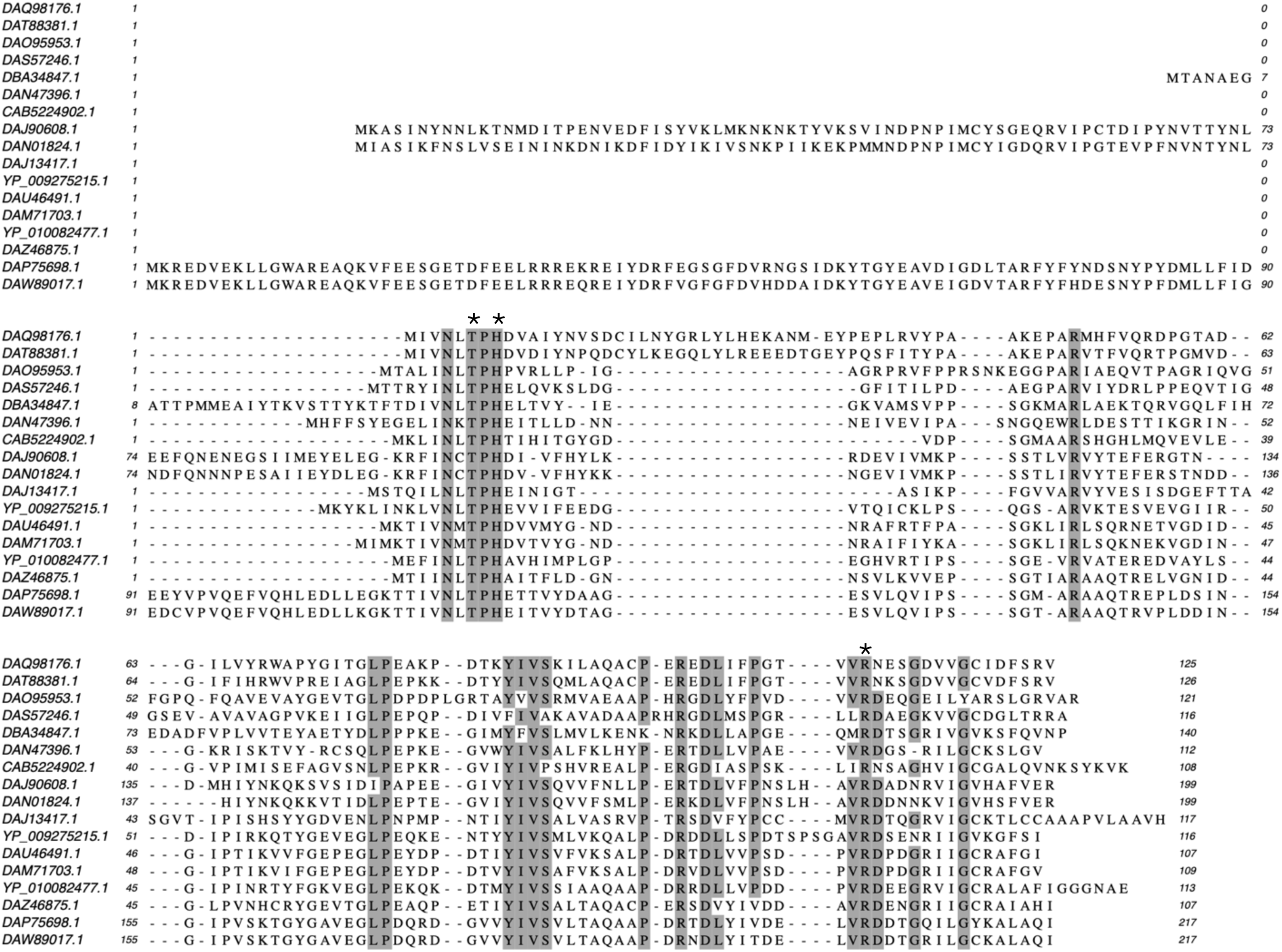
Sequence alignment of bacteriophage and plasmid Crn4 orthologues. T12, H15 and R112 (AprCrn4a numbering) are highlighted with asterisks. Some members of the family have an 80-100 aa N-terminal extension.

**Supplementary Figure 3.**
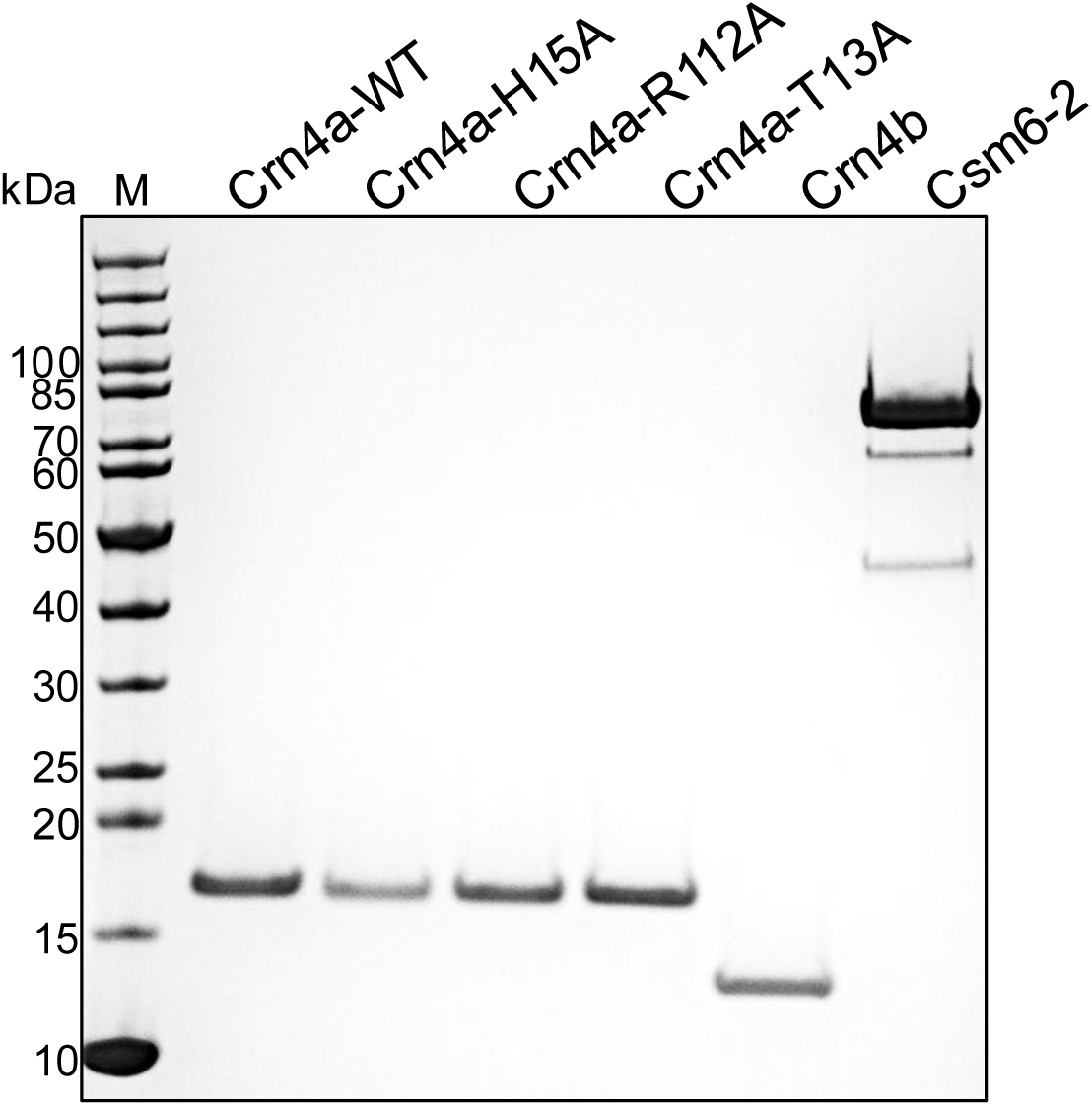
SDS-PAGE analysis of purified Crn4a, Crn4b and Csm6-2 proteins. The observed mass of Crn4a, Crn4b, and Csm6-2 on the gel is about 16, 14, and 86 kDa respectively, consistent with their theoretical mass. M is the molecular weight marker with sizes indicated.

**Supplementary Figure 4.**
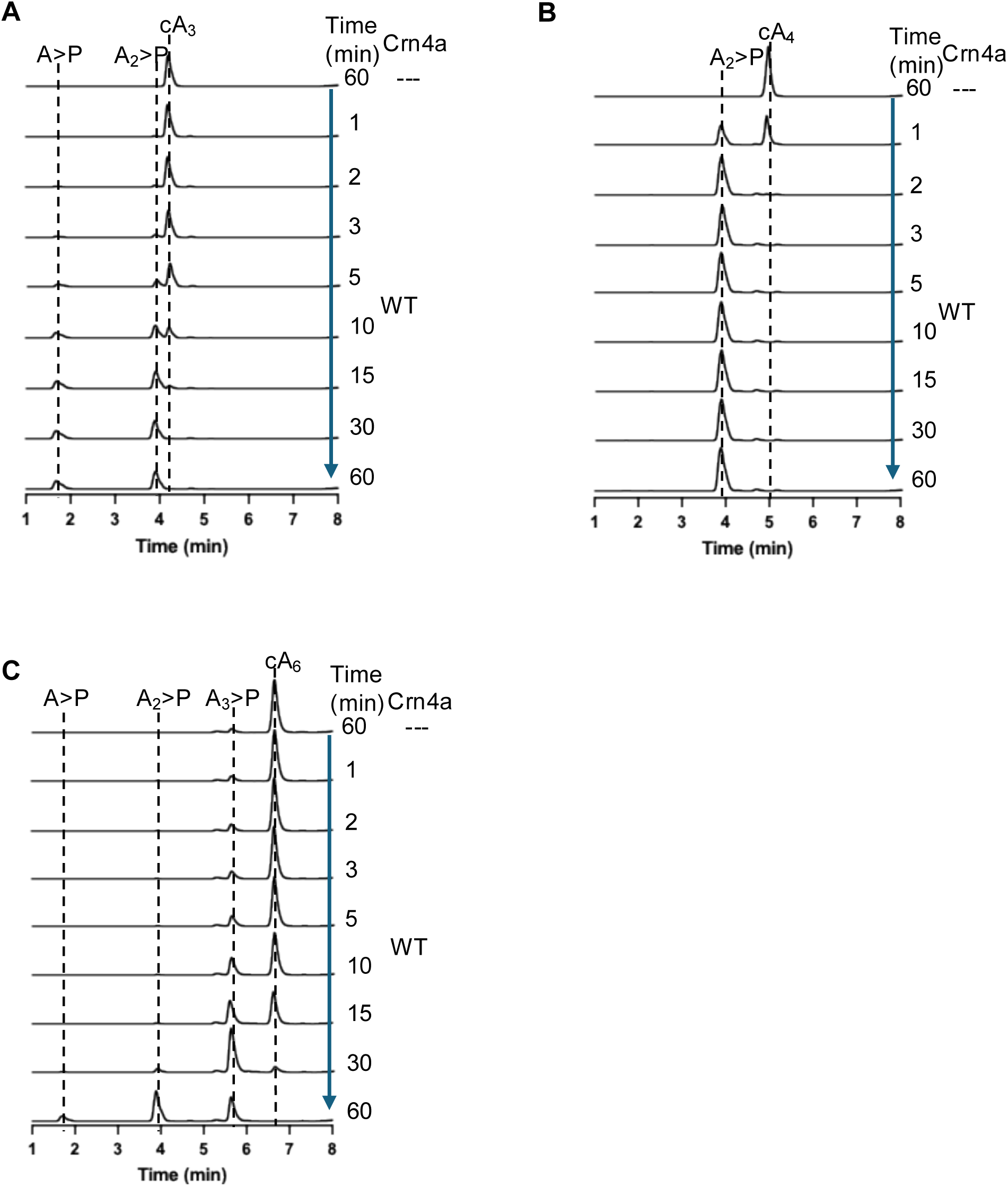
Time course of Crn4a ring nuclease activity. **A.** HPLC analysis of cA_3_-cleavage activity of Crn4a wild-type. 0.5 *μ*M Crn4a is incubated with 100 *μ*M cA_3_, cA_4_ in **B** and cA_6_ in **C** at 37 °C for the time points 1, 2, 3, 5, 10, 15, 30 and 60 min. Representative HPLC traces of three replicates are present.

**Supplementary Figure 5.**
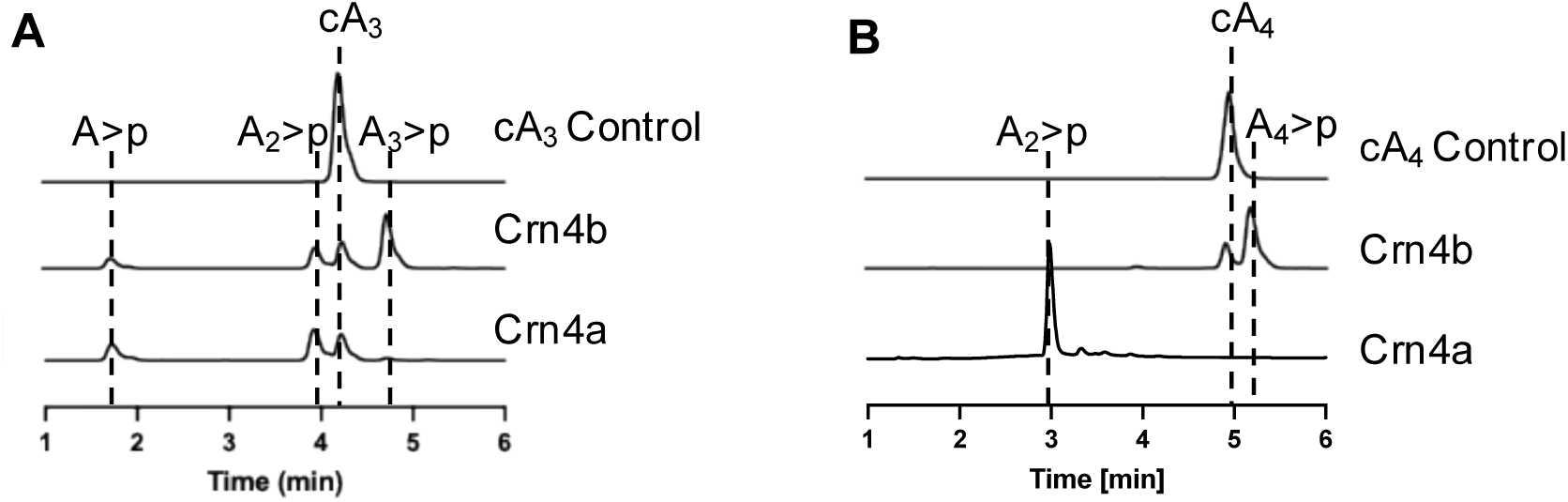
Comparison of ring nuclease activity of Crn4a and Crn4b. **A.** HPLC analysis of cA_3_-cleavage activity of Crn4a and Crn4b wild-type. 0.5 *μ*M Crn4 is incubated with 100 *μ*M cA_3_ (cA_4_ in **B)** at 37 °C for 10 min.

**Supplementary Figure 6.**
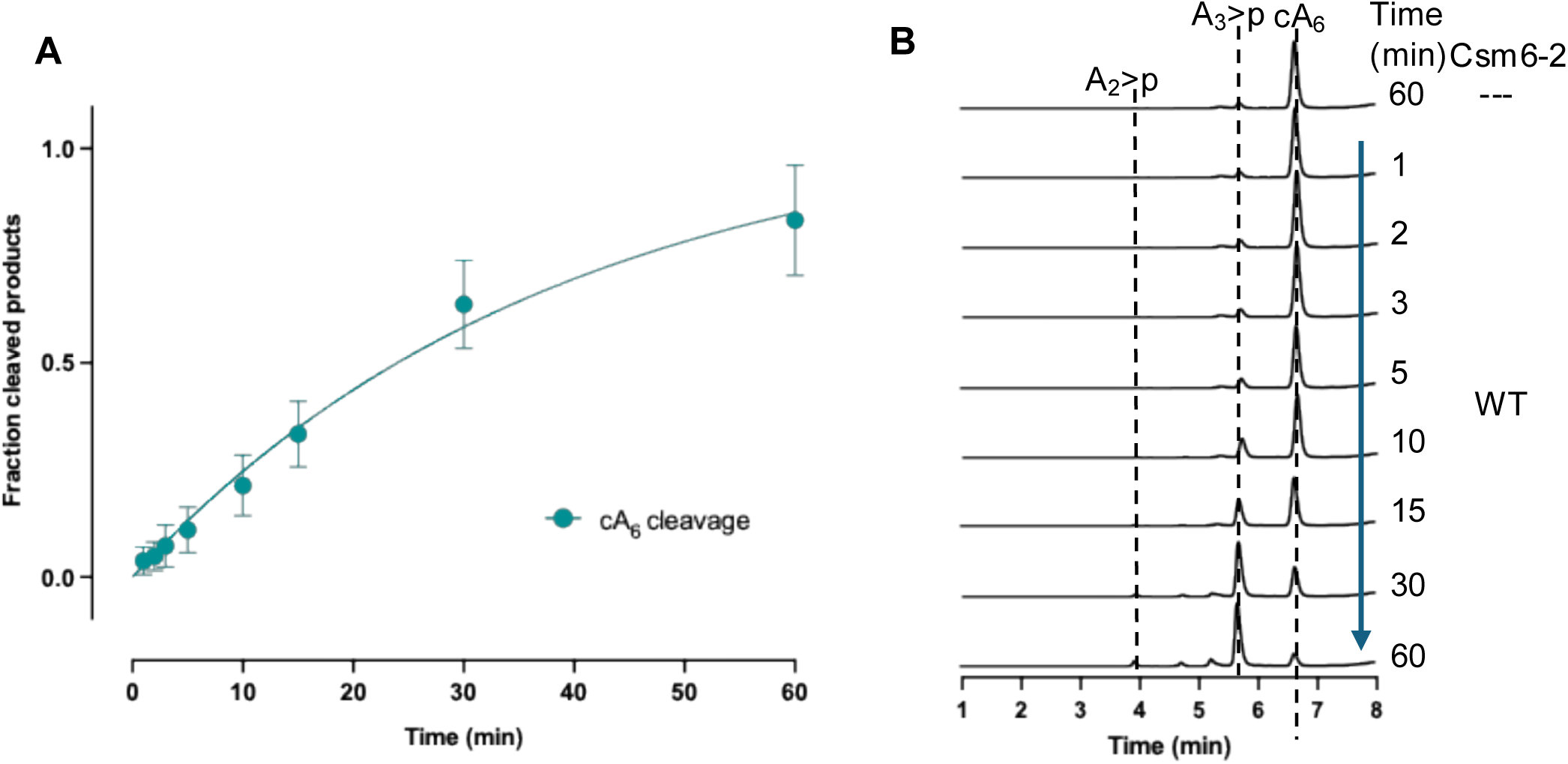
Csm6-2 ring nuclease activity against cA_6_. **A.** Kinetic analysis of cA_6_ cleavage activity. HPLC peaks of substrate and products were quantified and data plotted as fraction cleaved against time. Data points are shown as the means of triplicate experiments and standard deviation is shown. **B.** Representative HPLC traces of triplicate time course experiments are present.

**Supplementary Figure 7.**
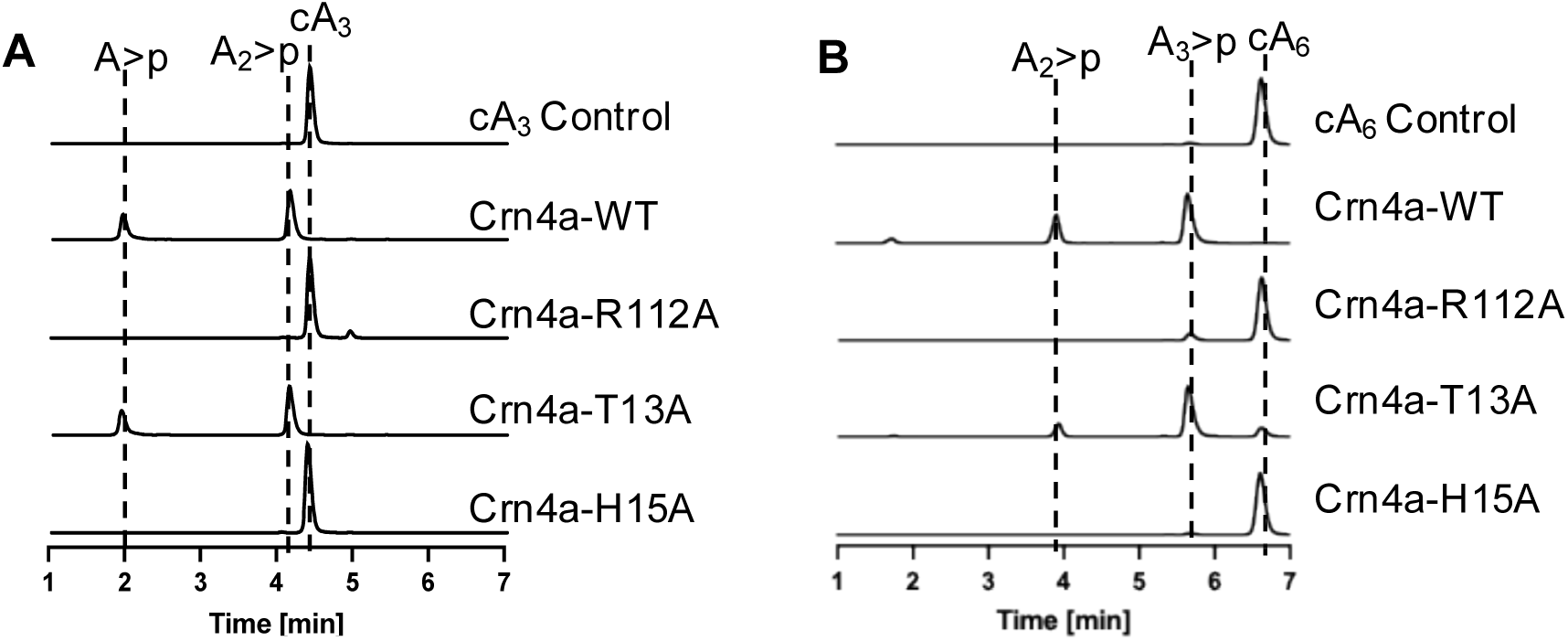
Ring nuclease activity of Crn4a WT and variants. HPLC analysis of cA_3_ cleavage activity (**A**) and cA_6_ cleavage activity (**B**) of Crn4a wild-type and variants. 0.5 *μ*M enzyme is incubated with 100 *μ*M cA_3_ or cA_6_ at 37 °C for 60 min.

**Supplementary Figure 8.**
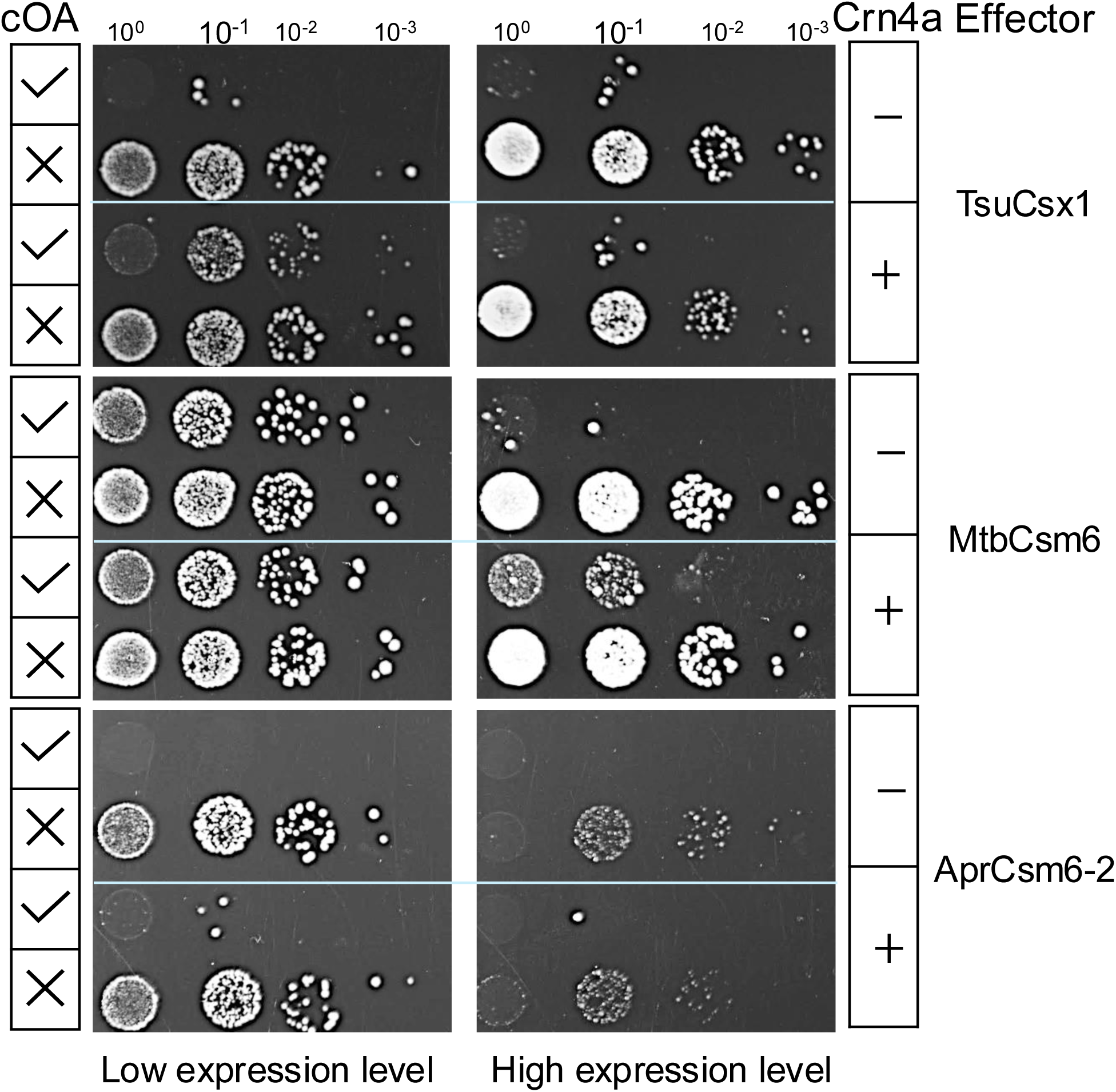
Comparison of plasmid immunity in low and high expression levels. C43 cells expressing MtbCsm system programmed with or without cOA production were transformed with plasmids harbouring effectors in the absence or presence of Crn4a. The transformants were selected on LB agar containing 0.2% (w/v) β-lactose and L- arabinose representative as a high expression level, while omitting both inducer presents as a low expression level. The pBAD and T7 promoter used in this systems were always observed a low level of transcription even in the absence of inducer.

**Supplementary Table 1.**
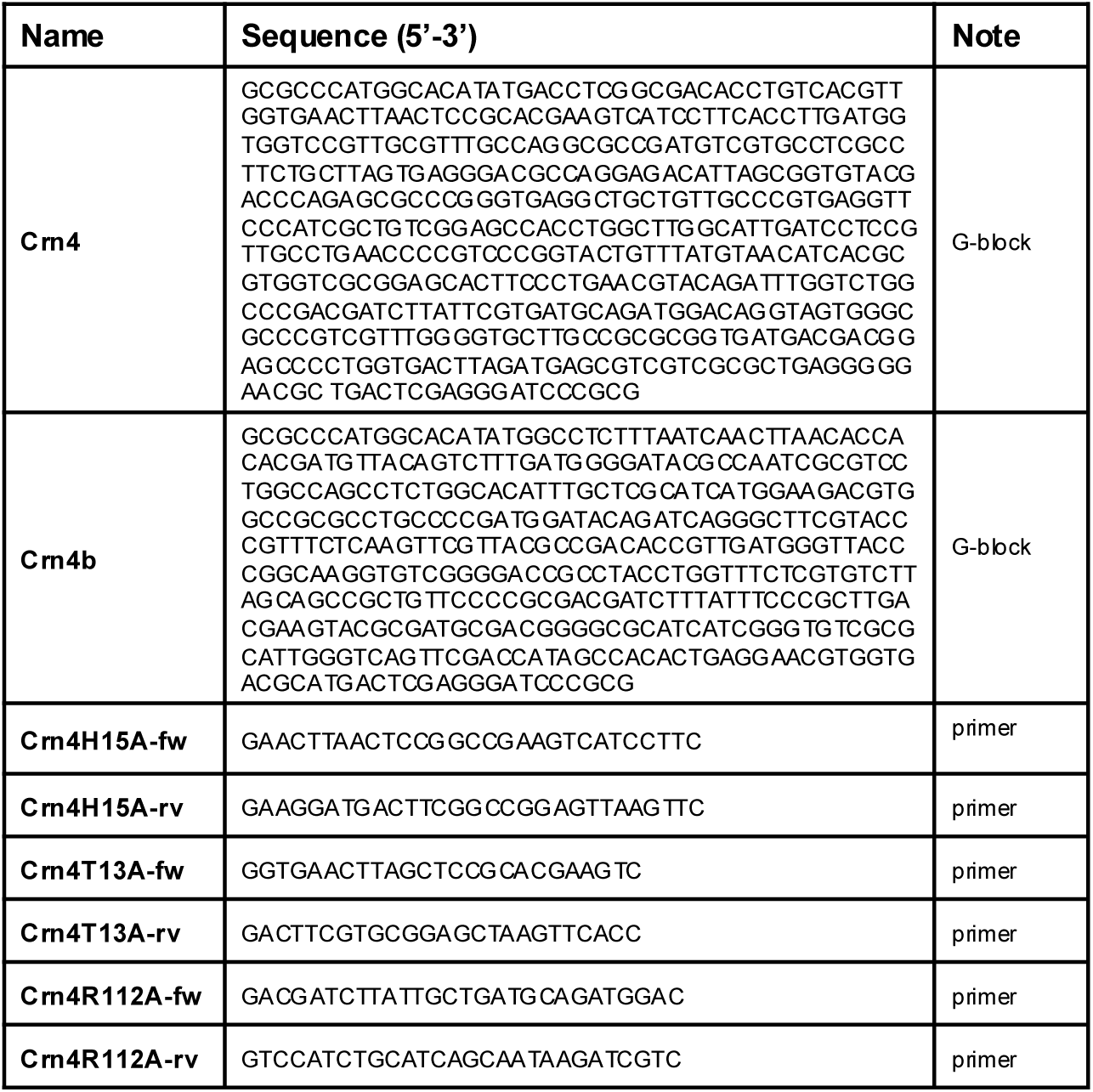
Synthetic genes and mutagenesis primers for Crn4 and Crn4b.

**Supplementary Table 2:** Data processing and refinement sta9s9cs for apo WT Crn4b and Crn4a H15A in complex with cA6.

|  | WT Crn4b | Crn4a H15A + cA<sub>6</sub> |
| --- | --- | --- |
| **Data processing** |  |  |
| Space group | C 1 2 1 | P 2<sub>1</sub> |
| Cell dimensions |  |  |
| a, b, c (Å) | 46.3, 61.3, 42.7 | 66.3 102.9, 72.4 |
| α, β, γ (°) | 90, 101.9, 90 | 90, 103.9, 90 |
| Resolution (Å) | 30.68 – 1.09 (1.11 – 1.09)* | 102.9 – 1.44 (1.47– 1.44)* |
| R<sub>merge</sub> | 0.036 (4.83)* | 0.068 (2.10)* |
| I/s(I) | 26.0 (0.2)* | 11.2 (0.3)* |
| Completeness (%) | 79.4 (10.3)* | 100 (99.2)* |
| Average redundancy | 11.9 (2.6)* | 6.9 (6.7)* |
| CC<sub>1/2</sub> | 1.000 (0.228)* | 0.998 (0.299)* |
| V<sub>m</sub> (Å<sup>3</sup>/Da) | 2.18 | 2.50 |
| Solvent (%) | 43.7 | 50.8 |
| **Refinement** |  |  |
| Unique reflections | 38596 (250) | 169690 (8333) |
| R<sub>work</sub> / R<sub>free</sub> | 16.8 / 18.5 | 22.2 / 24.2 |
| Geometric deviations |  |  |
| Bonds (Å) / Angles (°) | 0.005 / 1.295 | 0.004 / 1.164 |
| No. atoms (non H) |  |  |
| Protein | 896 | 5761 |
| Water | 98 | 694 |
| cA<sub>6</sub> | 0 | 396 |
| Mg<sup>2+</sup> | 0 | 16 |
| B factor (Å<sup>2</sup>) |  |  |
| Protein | 16.0 | 32.1 |
| Water | 29.9 | 36.2 |
| cA<sub>6</sub> |  | 20.8 |
| Mg<sup>2+</sup> |  | 27.1 |
| Ramachandran |  |  |
| Favoured / outlier (%) | 99.1 / 0 | 98.4 / 0 |
| Molprobity score / centile (%) | 1.01 / 98 | 0.93 / 100 |

\* Values in parentheses are for the high resolution shell.

